# Bacteriophage Treatment against Carbapenem-resistant *Klebsiella pneumoniae* (KPC) in a Neutropenic Murine Model of Gastrointestinal Translocation and Renal infection

**DOI:** 10.1101/2024.06.21.600121

**Authors:** Panagiotis Zagaliotis, Jordyn Michalik-Provasek, Eleftheria Mavridou, Ethan Naing, Ioannis S. Vizirianakis, Dimitrios Chatzidimitriou, Jason J. Gill, Thomas J. Walsh

## Abstract

**Background:** Carbapenemase producing *Klebsiella pneumoniae* (KPC) are globally emerging pathogens which that cause life-threatening infections. Novel treatment alternatives are urgently needed.

**Methods:** We therefore investigated the effectiveness of three novel bacteriophages (Spivey, Pharr and Soft) in a neutropenic murine model of KPC gastrointestinal colonization, translocation, and disseminated infection. Bacteriophage efficacy was determined by residual bacterial burden of KPC in kidneys. Parallel studies were conducted of bacteriophage pharmacokinetics and resistance..

**Results:** Treatment of mice with 5×10^9^ PFU of phage cocktail via intraperitoneal injection was effective in significantly reducing renal KPC burden by 10^2^ CFU (p<0.01) when administered every 24 hours and 10^3^ CFU (p<0.01) every 12 hours. Moreover, a combination of bacteriophage and ceftazidime-avibactam produced a synergistic effect, resulting in a 10^5^ reduction in bacterial burden in caecum and kidney (p<0.001 in both tissues). Prophylactic administration of bacteriophages via oral gavage did not prevent KPC translocation to the kidneys. Bacteriophage decay determined by linear regression of the ln of mean concentrations demonstrated R^2^ values in plasma of 0.941, kidney 0.976, and caecum 0.918, with half-lives of 2.5h < t_1/2_ < 3.5 h. Furthermore, a phage-resistant mutant displayed increased sensitivity to serum killing *in vitro*, but did not show significant defects in renal infection *in vivo*.

**Conclusions:** A combination of bacteriophages demonstrated significant efficacy alone and synergy with ceftazidime/avibactam in treatment of experimental disseminated KPC infection in neutropenic mice.

## INTRODUCTION

Carbapenemase-producing *Klebsiella pneumoniae* (KPC) is an emerging, highly-antibiotic-resistant pathogen, which is responsible for nosocomial infections worldwide(1,2). KPC encodes the *bla_KPC_* gene, which produces the carbapenemases. *Bla_KPC_* is located within a Tn3-type transposon, and is therefore easily transferred horizontally, thus spreading resistance within the species, as well as from one geographic location to another(3). Efforts to address these clinical challenges include enhanced infection control practices, better screening methods, optimal usage of existing antibiotics, and development of novel antimicrobial agents(4).

The *bla_KPC_* gene is the most prevalent cause for carbapenemase production. It first appeared approximately 30 years ago, in 1996, in the United States, and has since become endemic in large parts of the world, among which are New York, Greece, and Israel. Specifically, the sequence type (ST) 258, which was studied herein, encoding KPC-2, constitutes approximately 70% of all KPC isolates(1).

KPC mainly colonizes the alimentary tract, and many epidemiological studies have shown that most types of KPC infection are preceded by gastrointestinal colonization(5). KPC translocates through a disrupted gastrointestinal epithelial barrier into the circulation and subsequently into deep tissues, including the kidneys, which is facilitated even further during neutropenia(6).

As the therapeutic alternatives for treatment of *K. pneumoniae* infections are becoming more limited due to the rapid dissemination of resistant phenotypes worldwide, novel therapeutic strategies are urgently needed. One such strategy is the use of bacteriophages, which are bacteria-killing viruses, first discovered in 1917(7). Although the use of bacteriophages was largely abandoned in the second half of the 20th century due to the production of antibiotics, they have received increased attention as promising antimicrobial agents within the last three decades. Numerous preclinical studies have illustrated the efficacy of bacteriophages against different pathogens(8), and in addition, bacteriophages have been used in clinical settings alone or in combination with antibiotics, to treat refractory bacterial infections with a relatively high success rate(9–12).

In this study, we investigated the efficacy of bacteriophage treatment in a neutropenic murine model of KPC gastrointestinal translocation on the residual bacterial burden in renal infection (Fig. 1). Bacteriophage treatment when applied alone demonstrated significant reduction in residual bacterial burden. When used in combination with ceftazidime/avibactam, bacteriophage treatment produced a synergistic eradication of renal infection. This efficacy of bacteriophage treatment establishes a preclinical foundation for further investigation of this modality in neutropenic hosts with serious KPC infections.

**Figure 1.**
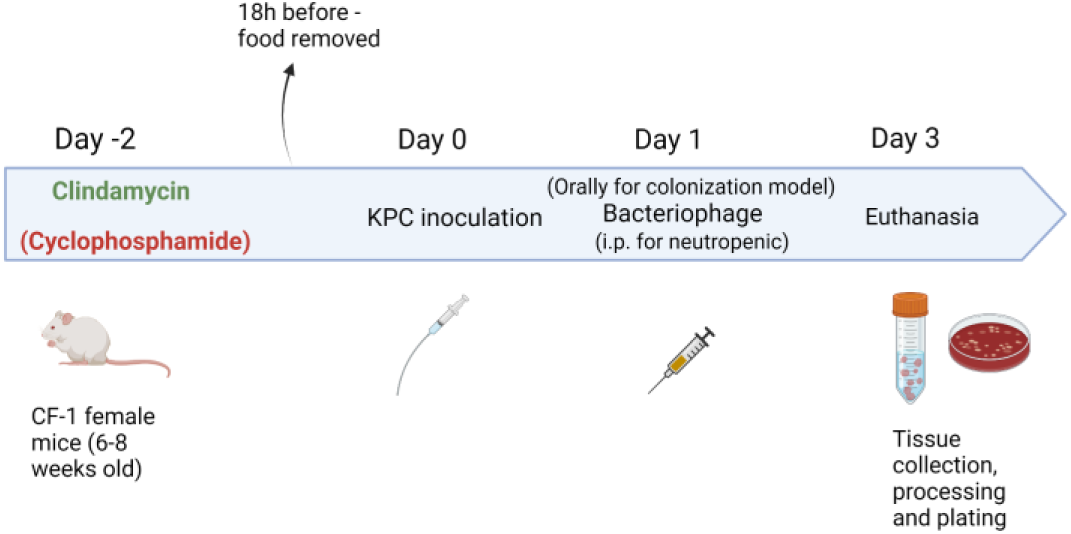
– Experimental design for inoculation of neutropenic mice with KPC

## RESULTS

### Efficacy of bacteriophage treatment on KPC burden in kidneys of neutropenic mice

Bacteriophage treatment resulted in an approximate 2 log_10_ reduction in bacterial burden in the kidneys of the mice when administered every 24 hours (p<0.01), and in an approximate 3 log_10_ reduction when administered every 12 hours (p<0.01), in comparison with the untreated control groups (**Fig. 2**). Bacterial survivors collected from these experiments (n=118) indicated that the majority of bacteria surviving phage treatment remained sensitive to phages Pharr and Soft (**Table 1**). Resistance to phage Spivey, which is only able to infect Pharr- and Soft-resistant mutants, was not observed.

**Figure 2.**
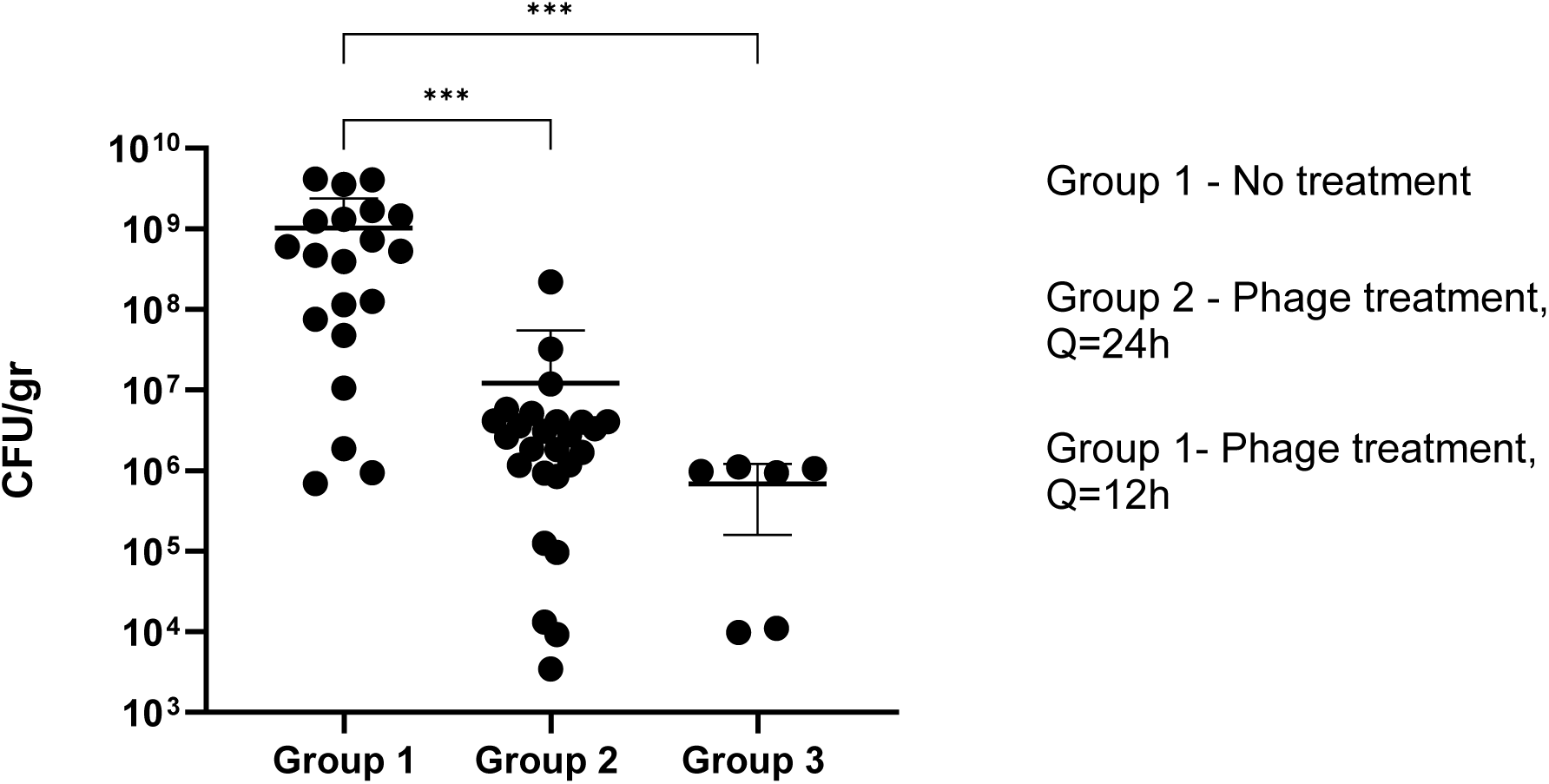
– Efficacy of bacteriophage treatment on KPC burden in kidneys of neutropenic mice. Residual bacterial burden (CFU/gr) in the kidneys of untreated controls (group 1) and groups treated with phage every 24 hours (group 2) and every 12 hours (group 3). When treated with 5×10^9^ PFU of the phage cocktail via i.p. injection every 24 hours (group 2), a 10^2^ CFU reduction was observed (p=0.001***); whereas, a 10^3^ CFU reduction was recorded for animals treated with the same concentration of the phage cocktail i.p. every 12 hours (group 3) (p=0.001***).

**Table 1.** *In vitro* susceptibility of bacterial colonies recovered following *in viv*o phage treatment to phages. Bacteria resistant to all three phages were not recovered.

|  | Colonies Tested | Pharr & Soft Susceptible | Percent Pharr & Soft Susceptible |
| --- | --- | --- | --- |
| Q8 Kidney | 9 | 9 | 100% |
| Q8 Cecum | 16 | 14 | 88% |
| Q12 Kidney | 25 | 21 | 84% |
| Q12 Cecum | 27 | 19 | 70% |
| Q24 Kidney | 21 | 15 | 71% |
| Q24 Cecum | 20 | 15 | 75% |
| Total | 118 | 93 | 79% |
| Kidney Only | 55 | 45 | 82% |
| Cecum Only | 63 | 48 | 76% |

### Prevention of KPC translocation by bacteriophage

In order to study the effect of bacteriophage on KPC translocation, the phage cocktail was administered at different time points (12h, 8h, 6h and 2h) to mice prior to their inoculation with KPC. There was no evidence of bacteriophage preventing KPC translocation in the neutropenic mouse model (**Fig. 3**).

**Figure 3.**
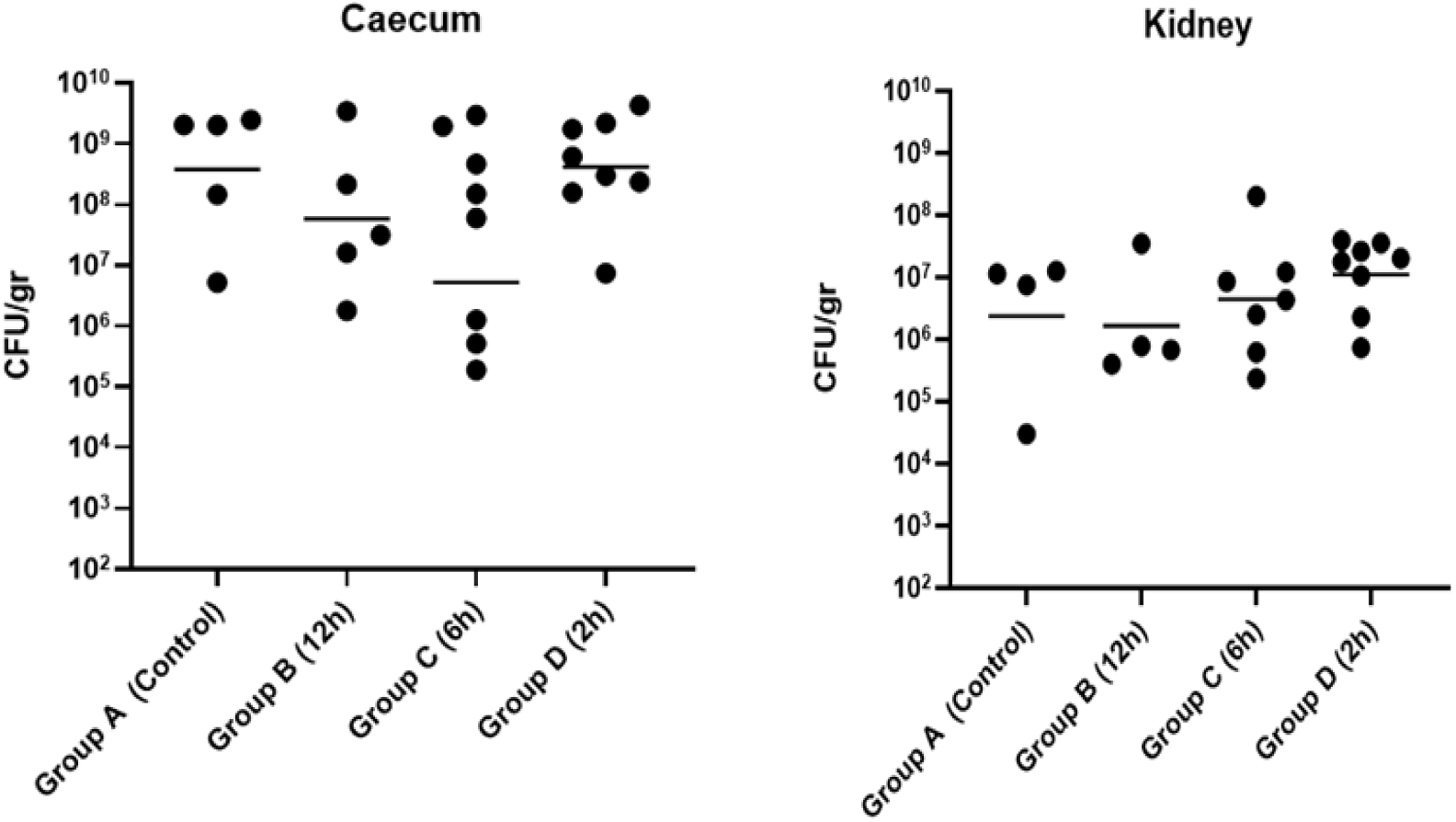
– Bacteriophage does not prevent KPC gastrointestinal translocation. **A**: Caecum CFU/g of pretreated groups with phage (12h, 6h and 2h prior to KPC inoculation) and untreated control groups (no phage administration). **B:** Kidney CFU/g of pretreated groups with phage (12h, 6h and 2h prior to KPC inoculation) and untreated control groups (no phage administration). Bacteriophage treatment prior to inoculation with KPC did not result in a significant reduction in bacterial burden in caecum, and did not prevent translocation to the kidneys.

Efficacy of simultaneous administration of bacteriophage and ceftazidime/avibactam on KPC burden in kidneys of neutropenic mice.

In a study of bacteriophage and CAZ/AVI simultaneously administered to persistently neutropenic mice, the combination resulted in a synergistic reduction of bacterial burden in the kidney (p<0.001) (**Fig. 4**). Groups 2, 4 and 5, which only received one form of treatment (either CAZ/AVI only or phage only) also showed significant reduction in bacterial burden (p<0.01).

**Figure 4.**
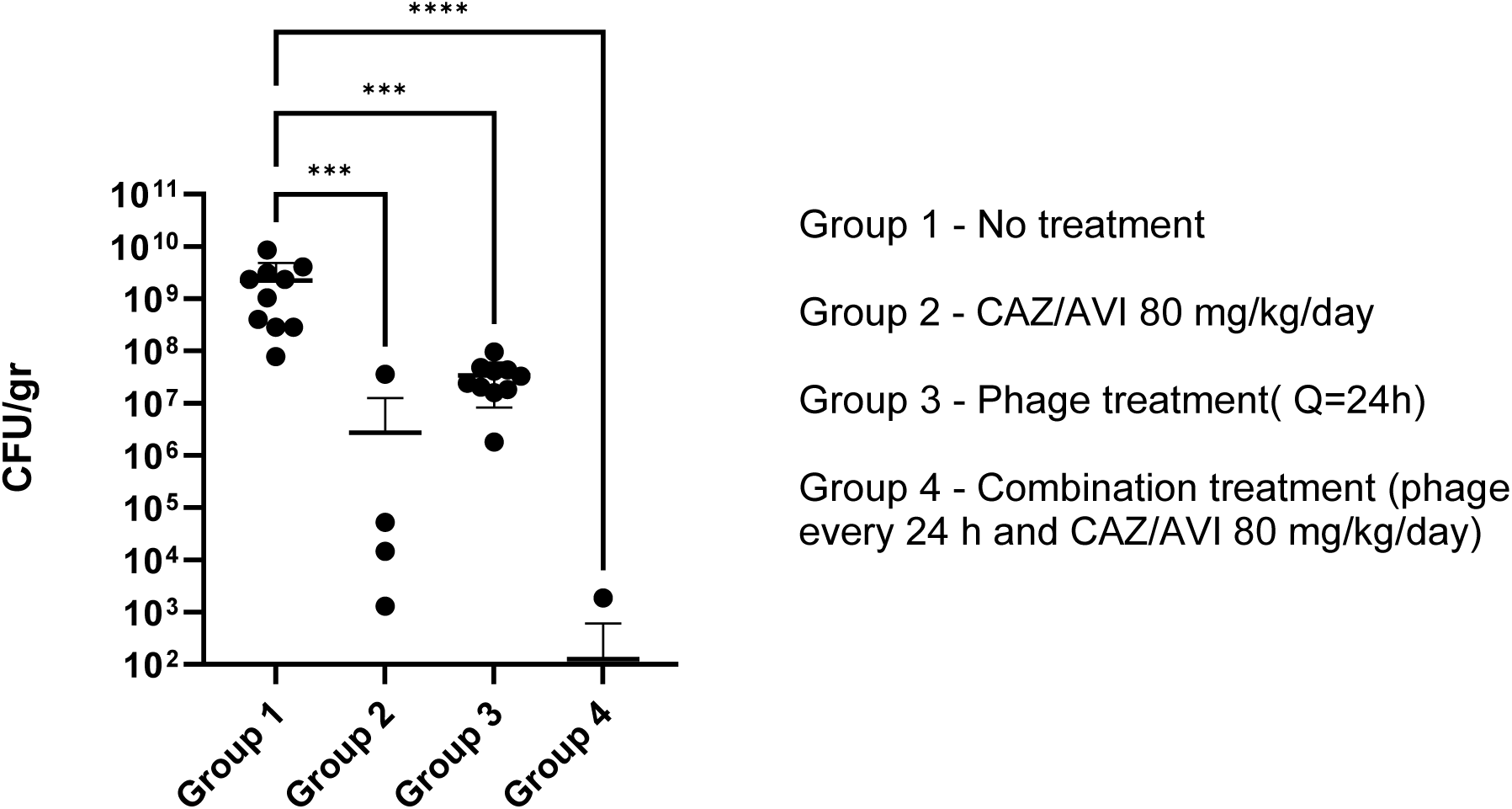
– Efficacy of simultaneous administration of bacteriophage and ceftazidime/avibactam on KPC burden in kidneys of neutropenic mice. The combination of bacteriophage and CAZ/AVI resulted in synergistic reduction of KPC kidney burden (CFU/gr) (****p<0.001). Treatment with phage only and CAZ/AVI only produced an approximate 2 log_10_ and 2.5 log_10_ reduction, respectively, in KPC kidney burden (***p<0.01).

### Phage-antibiotic interactions *in vitro*

The combination of caz/avi and phage did not appear to be antagonistic *in vitro*, but an enhanced antimicrobial effect between phage and caz/avi treatment was not observed (**Fig. 5**). Phage-antibiotic synergism was observed in our previous study and in other studies (13,14) but the mechanisms of this interaction remain unclear. Antibiotic sensitivity was found to be largely unchanged between the parental *K. pneumoniae* strain 39427 and its Pharr-resistant derivative (**Table 2**), indicating that the increase of phage resistance is not linked to increased antibiotic sensitivity as has been observed in other systems (15,16).

**Figure 5.**
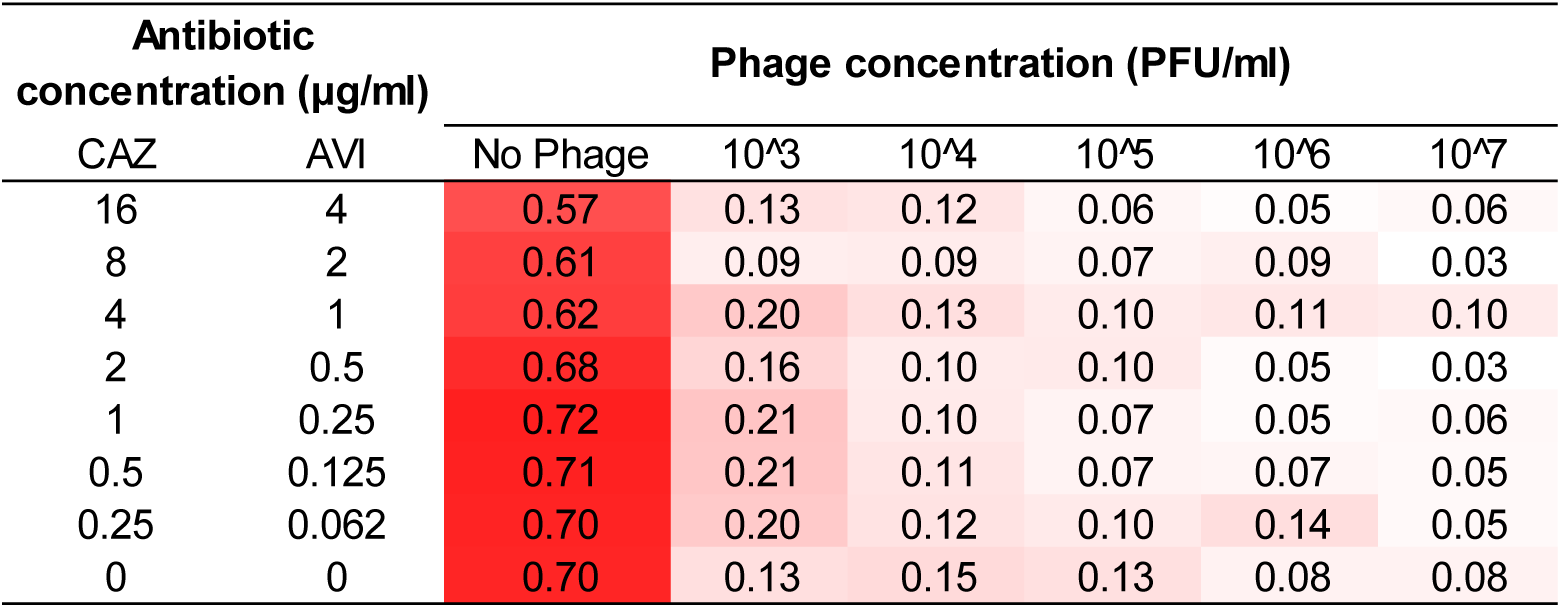
Endpoint checkerboard assay incorporating the triple phage cocktail and ceftazidime/avibactam. The activity of bacteriophage alone (bottom row of the table) markedly reduced the optical density of KPC suspension by as much as 90%.

**Table 2:** Minimum inhibitory concentration assays of the wild-type KPC strain 39427 and its Pharr-resistant derivative using commercial ESBL broth microdilution assays. The phage-resistant mutant was not observed to be more sensitive to antibiotics than its parental strain.

| Antibiotic | Minimum inhibitory concentration |  |
| --- | --- | --- |
|  | 39427 wt | 39427 Pharr-R |
| Ceftriaxone | >128 | >128 |
| Cephalothin | >16 | >16 |
| Cefotaxime | >64 | >64 |
| Cefotaxime/clavulanic acid | 64/4 | 64/4 |
| Ceftazidime | >128 | >128 |
| Cefazidime/clavulanic acid | >128/4 | >128/4 |
| Imipenem | >16 | >16 |
| Cefepime | >16 | >16 |
| Cefpodoxime | >32 | >32 |
| Cefoxitin | >64 | >64 |
| Piperacillin/tazobactam | >64/4 | >64/4 |
| Meropenem | >8 | >8 |
| Gentamicin | 4 | 4 |
| Ciprofloxacin | >2 | >2 |
| Ampicillin | >16 | >16 |
| Cefazolin | >16 | >16 |

### Bacteriophage kinetics in plasma, caecum and kidney

After collecting plasma, caecum and kidney from animals at 2, 4, 6, 8, 12, and 24 hours post-inoculation with KPC, bacteriophage titers were measured with the spot titer method for plasma and kidney, and with the full-plate titering method for caecum. Measurements showed that bacteriophage titers were reduced in a linear manner from all three sample types with a half-life ranging from 2.5 to 3.5 hours (**Fig. 6**).

**Figure 6.**
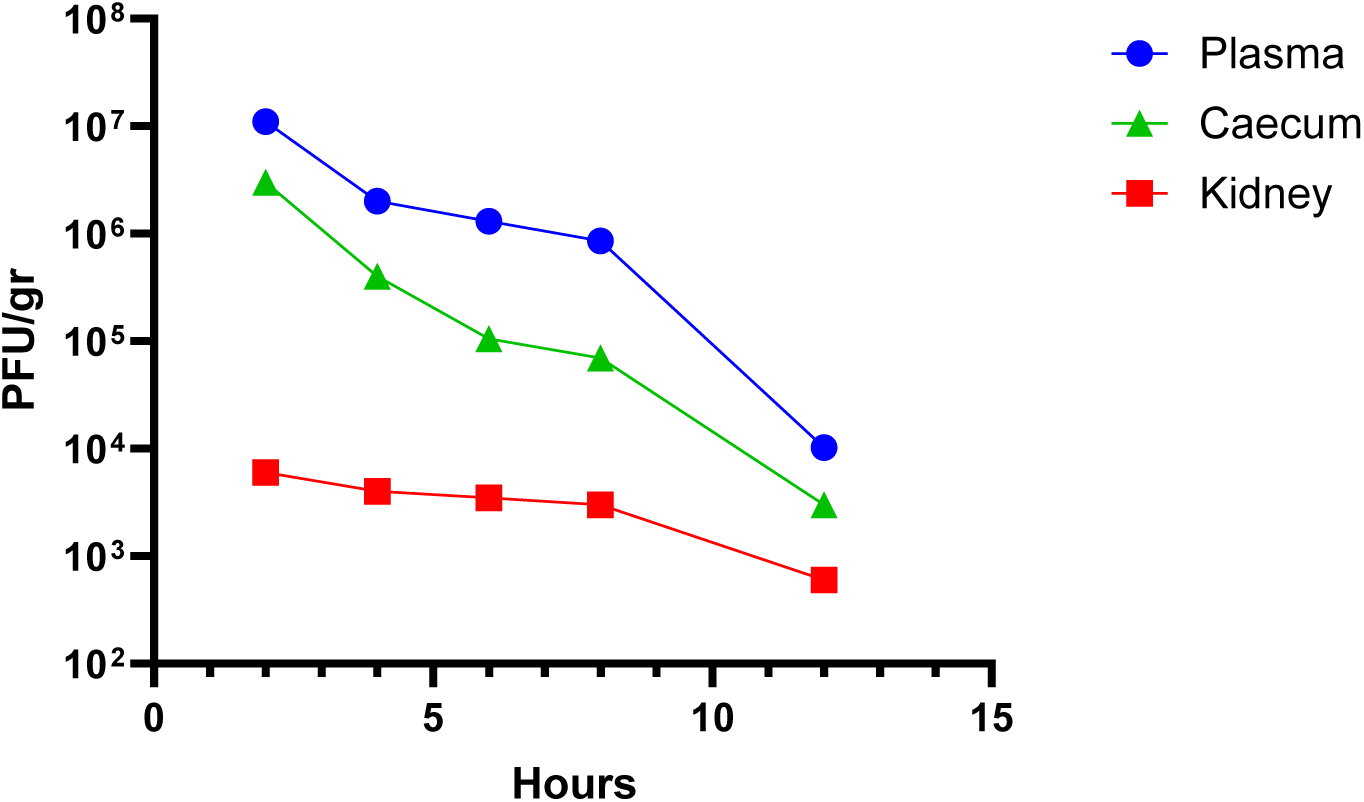
Bacteriophage kinetics in plasma, caecum and tissue. Bacteriophage concentration in plasma, caecum and kidney after a single 100μL i.p. dose of 5×10^10^PFU phage lysate. Titers were reduced linearly on the log scale in plasma, caecum and tissue. The half-life ranged from 2.5 hours to 3.5 hours for plasma and cecum, while the half-life in kidney tissue was approximately 6 hours.

### Susceptibility of phage-resistant mutans to complement system killing *in vitro*

In order to further understand the effect of phage resistance of KPC on virulence, in an *in vitro* serum killing assay was performed to determine the effect of complement on a phage Pharr-resistant mutant. This mutant, which is missing its capsule (17,18), demonstrated increased susceptibility to complement killing by human serum (**Fig. 7**).

**Figure 7:**
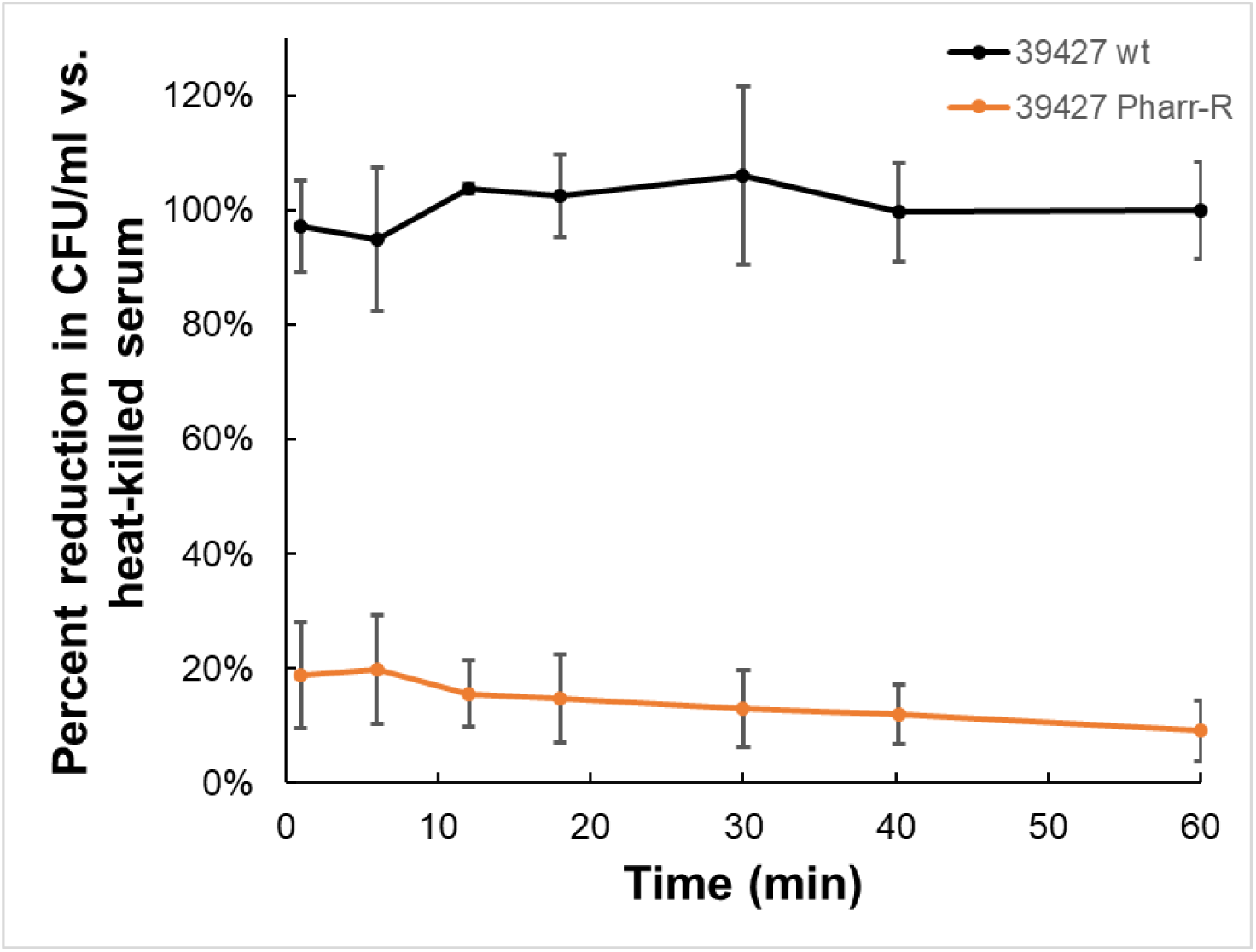
Bacteriophage Resistant KPC Mutants are Complement Susceptible. KPN39427 WT, and the Pharr-resistant bacterial mutants were treated with either competent human serum or heat killed (HK) human serum and assessed for bacterial survival over a 60 minute period. Data shown is adjusted to the number of surviving cell of the same strain incubated in heat-hilled serum. A mean of 3 biological replicates is represented, error bars indicate standard deviation. The wild-type strain 39427 is not affected by complement killing, while the Pharr-resistant mutant is sensitive to killing by complement.

### Effects of phage resistance on bacterial fitness *in vivo*

Structural changes in some strains of KPC that are resistant to bacteriophages may result in a loss of fitness. We therefore examined whether the loss of capsule observed in the Pharr-resistant mutant of KPC affected the ability of the bacteria to colonize mouse tissues. Neutropenic mice, premedicated with clindamycin and cyclophosphamide following the protocol for KPC translocation, were inoculated via oral gavage with either 10^7^ CFU of the wild type KPC or the Pharr-resistant mutant. The ability of the Pharr-resistant mutant of KPC to colonize the caecum, translocate and infect the kidney, was not significantly affected by the loss of capsule. This mutant displayed residual CFUs within the renal tissue that were comparable to those of the wild type KPC (**Fig. 8**).

**Figure 8:**
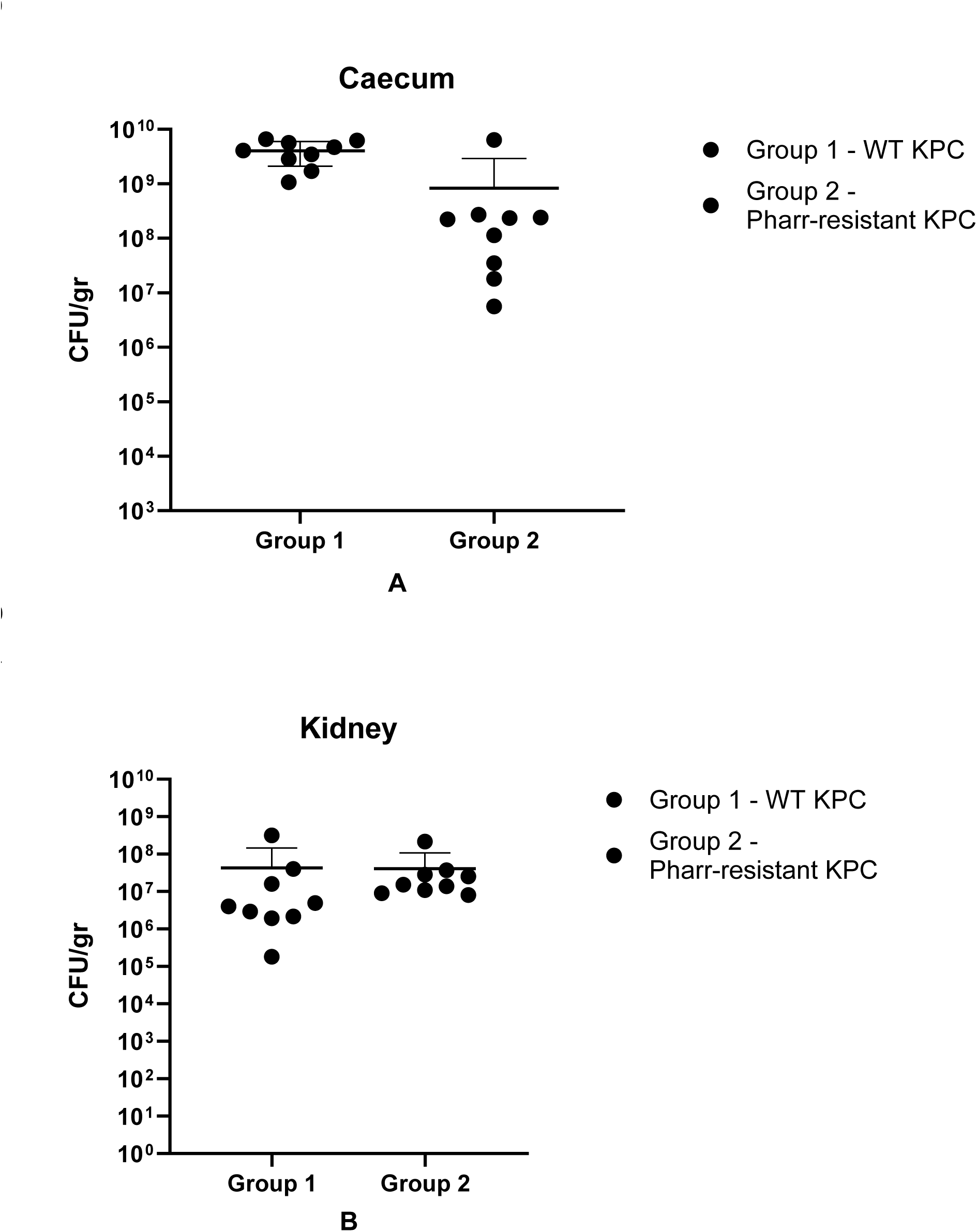
Effects of phage resistance to bacterial virulence *in vivo*. **A**: Caecum CFU/g, indicative of the colonization ability (and therefore the virulence) of phage-resistant mutants derived from phage-treated groups, in comparison with the WT. **B:** Kidney CFU/gr, indicative of the extent of translocation (and therefore the virulence) of phage-resistant mutants derived from phage-treated groups, in comparison with the WT.

## DISCUSSION

The results of this work highlight the efficacy of a bacteriophage cocktail comprised of three phages against KPC renal infection in a neutropenic murine model. In addition to its efficacy against KPC colonization, this phage cocktail was shown to reduce KPC burden in kidneys by approximately 2 log_10_ (p<0.01) when administered every 24 hours (Fig. 2). This effect was pharmacodynamically enhanced by more frequent administration of the phage cocktail (every 12 hours), resulting in a 3 log_10_ reduction of renal KPC burden (p<0.01). Despite its marked effectiveness against renal infection, prophylactic administration of the phage cocktail was not shown to be efficient in preventing KPC translocation in neutropenic murine hosts (Fig. 3). Notably, even a small residual amount of KPC in the cecum was sufficient to result in renal translocation in neutropenic hosts.

As most patients with KPC bacteremia will be receiving an antimicrobial agent (4), understanding the potential for antagonistic or synergistic interaction between CAZ/AVI and phage is important. The *in vivo* data presented herein illustrated a significant reduction in renal KPC burden (nearly 5 log_10_) following treatment by a combination of phage and antibiotic (Fig. 4). The mechanism for this synergistic activity is not clear, but an additional effect of this action is the elimination of potential resistance to either one of the agents, which is a phenomenon commonly occurring when using a single-agent treatment approach (19). To better understand the protective phenotype seen in the *in vivo* experiments where mice were best protected by a treatment of both bacteriophage and Ceftazidime/Avibactam (CAZ/AVI) we attempted to replicate this synergy *in vitro* with a plate assay. This assay was designed using the effective concentrations observed in the MIC experiments, as well as previous bacteriophage growth inhibition assays. In general, the combination of CAZ/AVI and phage was not synergistic *in vitro*, but possibly additive (Fig. 5). The aggregate *in vitro* and *in vivo* data indicate that bacteriophage is compatible with a standard-of-care treatment, and that there is significant benefit to treatment of KPC infection with both bacteriophage and CAZ/AVI. The recent emergence of resistance of KPC strains to CAZ/AVI further underscores the importance of combination therapy with bacteriophage (20,21).

Bacteria are in a constant evolving race to face the challenges posed to them by their natural predators. As a result, bacteriophage resistance is a phenomenon which can potentially occur during the course of bacteriophage treatment(22). A common hypothesis governing past years of bacteriophage research proposed that phage-resistant mutants which would result from treatment would display increased susceptibility to antibiotics(17,18). The mechanism for this increased susceptibility to antibacterial agents is attributed to alterations in their outer cell membrane and/or capsule, which would permit higher concentrations of antibiotics to enter the bacterial cell and to exert their activity(23). In this work, defects in capsule synthesis are known to confer resistance to phage Pharr (17,18) and by extension to phage Soft, which is unable to infect Pharr-resistant mutants. However, the majority of the tested bacterial clones collected from phage-treated animals showed that most were still sensitive to Pharr and Soft (**Table 1**), suggesting that bacterial avoidance of phage, such as in protected sites, was more of a limiting factor in treatment than was the emergence of resistance. When studying potential MIC changes of several antibiotics against the phage-resistant mutant, no changes in sensitivity were observed (**Table 2**), indicating that the occurrence of bacteriophage resistance does not necessarily result in increased susceptibility to conventional antibiotics. When further exploring the potential mechanism behind the observed *in vivo* synergy in an *in vitro* experiment of complement system killing in serum, it was observed that the phage-resistant mutant is susceptible to complement system killing (Fig. 7). This observation highlights a potential mechanism for loss of fitness *in vivo*, which would include reduced protection from complement killing due to the loss of capsule(24,25).

Our studies of bacteriophage kinetics in plasma, cecum, and kidney tissues of infected mice revealed a linear reduction on the logarithmic scale, with a half-life of 2.5-3.5 hours in plasma and cecum, and a half-life of approximately six hours for kidneys and with sufficient bacteriophage titers in renal tissue to potentially exert sustained therapeutic effect (Fig. 6).

In summary, this work highlights the efficacy of bacteriophage treatment, alone or in combination with antibiotics, against renal KPC infection in a neutropenic murine model and has important clinical implications for treatment of these lethal infections in neutropenic patients with the potential to be used alone or in combination with existing small molecule antimicrobial agents.

## MATERIALS AND METHODS

### Bacterial strains

The KPC strain 39427 [REF, NZ_CP054268] was kindly provided by Dr. Michael Satlin (Weill Cornell Medicine) and previously studied in our laboratory(26,27). This strain and its phage-resistant derivatives were used for all *in vitro* and *in vivo* experiments in the murine model. Bacteria were routinely cultured on tryptic soy broth (TSB, Difco) or tryptic soy agar (TSA, Difco) aerobically at 37°C. The minimum inhibitory concentration (MIC) of ceftazidime/avibactam (CAZ/AVI) for strain 39427 is 2μg/mL(26,27).

In preparation of the KPC inoculum, bacteria were prepared by suspending a bacterial colony from TSA agar into 50ml of TSB and shaking at 37°C for 4h. When an optical density of 0.01 at 600nm (OD_600_ = 0.01) was reached, corresponding to 10^7^ CFU/ml, the culture was centrifuged, and the resulting pellet was resuspended twice in sterile normal saline (0.9% NaCl). OD_600_ was measured again, for a resulting inoculum of 10^7^ CFU, which was administered to the animals via oral gavage.

### Bacteriophages

Bacteriophages Pharr (NC_048175), Soft (NC_048805) and Spivey (MK630230) were used in all experiments. Phages were routinely titered by the soft agar overlay method(28) using TSA bottom plates and top agar consisting of 10 g/L Bacto^™^ tryptone (VWR), 10 g/L NaCl and 0.5% Bacto^™^ agar (VWR), supplemented with 5 mM each of CaCl_2_ and MgSO_4_. Host lawns were established by inoculating 4 ml of molten top agar with 0.1 ml of a mid-log (OD_550_ 0.3-0.5) KPC host culture. For administration in murine models, phage lysates were prepared by inoculating log-phase, 1 L TSB liquid cultures of *K. pneumoniae* strain 1776pc (for phages Pharr and Soft) or 1776pcPharr-R (for phage Spivey) with phage at a multiplicity of infection (MOI) of 0.001-0.1 and culturing until lysis. Phage lysates were cleared by centrifugation (8,000 x g, 4°C, 10 min), filter-sterilized, and purified by isopycnic CsCl gradient ultracentrifugation as described previously(29). Residual endotoxin was removed by passage through an Endotrap HD column (Lionex), filter-sterilized, and final endotoxin was determined to be less than 1 EU/ml by EndoZyme II recombinant factor C assay (bioMerieux).

### Animals

All animal studies described herein were approved by the Weill Cornell Research Animal Resource Center (RARC), following the guidelines of the Institutional Animal Care and Use Committee (IACUC – protocol number: 2015-0043). Female 6-8 week old CF-1 mice weighing 26-30 g (Charles River Laboratories) were used in all experiments. Mice at that age are considered adults, and their weight allowed for reproducible data and easier dosing calculations. Outbred mice were used, aiming for an extrapolation of the data to the genetic diversity that is observed in humans.

### KPC colonization and translocation in neutropenic mice

Female, 6-8 weeks old CF-1 mice, weighing 26-30g, were rendered neutropenic by administration of cyclophosphamide, and resident gastrointestinal bacterial microbiome was disrupted by administration of clindamycin prior to inoculation with KPC strain 39427 via oral gavage. The day of KPC inoculation was designated as day 0 of the experiment. Mice were administered 300 mg/kg cyclophosphamide IP on day -2, and 150 mg/kg cyclophosphamide IP on each of days 0 and 3 (Fig. 1). Starting on day -2, mice also received a loading dose of clindamycin 1.5 mg IP per mouse. Each subsequent day until the end of the experiment, they received clindamycin 1.5 mg/kg IP. Eighteen hours prior to KPC inoculation, food was removed from mouse enclosures to prevent coprophagy. On day 0, a KPC inoculum of 10^6^ CFU/mouse (0.1ml of a OD=0.01 culture, corresponding to 10^7^ CFU/ml) was administered to the mice via oral gavage, using a blunt 20-gauge oral gavage needle. On days 1 and 2 of the experiment, infection was confirmed by plating stool of the mice on HardyCHROM^™^ CRE agar plates (Hardy Diagnostics). Animals were monitored twice daily for humane endpoints, including hunched posture, squinting of the eyes, lethargy, ruffled fur, and weight loss. Whenever an animal exhibited such endpoints, it was euthanized and caecum, kidneys and plasma were collected for bacterial quantification and inflammatory biomarker measurement. On day 3 of the experiment, animals were euthanized by CO_2_ overdose, and their caecum and kidneys were collected in sterile normal saline (0.9 % NaCl). Blood was collected via cardiac puncture. The caeca and kidneys of the mice were weighed and homogenized using a tissue homogenizer (Omni). Tissue homogenates were serially diluted and plated on CRE selective agar plates (HardyCHROM) using the single plate - serial dilution spotting (SP-SDS) method(30). Plates were incubated at 37°C overnight, colonies with the expected dark blue CRE morphology were enumerated, and counts adjusted to CFU/g sample. The lower limit of detection of the assay was 10^2^ CFU/g.

### Prophylactic model of KPC translocation

In an effort to explore the possibility of prevention of KPC translocation with oral administration of bacteriophages, 5×10^9^ PFU of bacteriophage was administered via oral gavage to the mice, at three different time points: 12, 6, and 2 hours prior to inoculation with KPC. The mice were euthanized 24 hours after KPC inoculation, caeca and kidneys were collected as described previously, and CFU were determined as mentioned.

### Simultaneous administration of ceftazidime/avibactam (CAZ/AVI) and bacteriophage on bacterial burden in caecum and kidney

Animals were assigned to four study groups: The first group received no antibacterial treatment (untreated controls), the second group received CAZ/AVI (80 mg/kg/day IP divided in three doses), the third group received a combination of CAZ/AVI (80 mg/kg/day IP) and bacteriophage (5×10^9^ PFU/mouse), and the fourth group received bacteriophage-only treatment (5×10^9^ PFU). The experiment was conducted for four days, on which time the mice were euthanized with CO_2_, their caeca and kidneys collected, and CFU determined as previously described.

### Virulence studies

The effect of structural changes on the virulence of phage-resistant KPC mutants was examined by inoculating different groups of mice with each mutant and then performing quantitative cultures of KPC in kidneys and caeca. Group 1 was a wild-type infected group, and group 3 was inoculated with the Pharr-resistant mutant, lacking the O-antigen.

### Plasma sample preparation

Blood samples collected in EDTA tubes were centrifuged at 4°C, 4000 rpm for 15 minutes. Plasma was collected with a pipette, transferred to screw top 1.5 ml tubes (VWR^®^) and stored at -80°C.

### Serum bactericidal assay

KPC bacterial cultures diluted to 10^5^ CFU/ml in TSB and placed on ice were incubated in both serum and heat-killed serum at 1:10 ratio for final CFU estimated at 10^4^. Cells were incubated at 37°C with timepoints taken and plated on TSA plates for surviving CFU. The lower limit of detection was 50 CFU/ml.

### Bacteriophage susceptibility of KPC isolates recovered following *in vivo* treatment

Colonies cultured from murine kidney or cecal samples following multiple doses (Q24, Q12, Q8) of bacteriophage in the clindamycin and cyclophosphamide-immunosuppressed murine model of KPC infection were cultured and susceptibility to bacteriophages Pharr and Soft was conducted by spot titer assay.

### Antimicrobial susceptibility of WT and phage-resistant mutants

Bacterial susceptibility to antimicrobial agents was determined by microplate MIC assays using standard ESBL assay plates (Trek Diagnostics) and by custom-made plates designed to measure high ranges of antibiotic concentrations. Bacterial inoculum was standardized to between 1×10^5^ and 1×10^6^ CFU using a 0.5 McFarland turbidity standard and immediate serial dilution, with inoculated CFU/mL confirmed by plating to TSA plates. To each well, 100 μL of inoculum was added to each well of the ESBL plate, the plate was sealed, and incubated at 37°C for 18 hours before being read. Each well was given either a positive or negative score for growth, defined as the well having any growth being positive. From these positive/negative scores, the actual MIC was generated with the lowest concentration of antibiotic which inhibited bacterial growth being considered the MIC for that strain/antibiotic combination. The data were representative of two technical replicates.

### Checkerboard assay for *in vitro* evaluation of the efficacy of phage and CAZ/AVI combination

A checkerboard assay representing combinations of bacteriophage and CAZ/AVI was applied to determine growth inhibition of single agents and combinations following an 18h incubation. Combinations included the bacteriophage at 10^7^ to 10^3^ PFU or No Phage, and CAZ/AVI from 16 to 0.025 μg/mL or no antibiotic. Bacterial cultures were added at 5×10^5^ CFU final concentration as described previously. Final well volumes were 200 μl and plates were covered and incubated 18h at 37°C before results were read using a plate reader measuring absorbance at 550nm. Bacterial strains tested were 39427 WT, 39427 Pharr Resistant Mutant, 39427 EI resistant Mutant, and 39427 JR Resistant Mutant. These were all *in vitro* generated bacteriophage resistant bacterial mutants. Results shown represent 3 averaged replicates

## Acknowledgment

This work was supported by NIAID, 1R21/R33 AI121689-01. TJW also was supported as an Investigator of the Save Our Sick Kids Foundation.

